# 3D geometry and mechanics of a single apical stem cell ensure helically symmetric plant body in multicellular models

**DOI:** 10.64898/2026.02.24.707705

**Authors:** Naoya Kamamoto, Koichi Fujimoto

## Abstract

Self-renewing divisions of stem cells at the growing tip universally govern plant development. In basal land plants, a single apical stem cell (AC) undergoes successive 120-degree rotational divisions while maintaining a self-similar tetrahedral geometry with three division walls, recapitulating the helically symmetric body architecture. The mechanisms by which AC geometry and mechanics determine the division axis and its rotation are largely unknown. Here, we develop two 3D multicellular mathematical models to address these questions. Our geometrical model reveals that the least area rule, coordinated with the AC surface curvature, is sufficient to drive rotational divisions by rotating the geometric proportion of the AC. Furthermore, by incorporating biologically plausible cell growth mechanics, we show that the tetrahedral AC and its rotational division spontaneously emerge from a spherical cell. Notably, the maximal tension rule confers superior robustness against stochastic fluctuations in division orientation compared to the least area rule, because the sequential formation of division walls continuously updates the tension distribution to guide subsequent orientation. These results suggest that the tetrahedral AC and the helically symmetric body plan are natural consequences of self-renewing cell divisions, imposed by 3D geometry and/or mechanics.

## 1 Introduction

The evolutionary development of three-dimensional multicellular bodies is the pivotal event for land plants to stand out from their surroundings and flourish on the land (Harrison, 2017; Moody, 2020). Since plant cells cannot rearrange themselves, precise orientation of cell division is fundamental for 3D development (Smolarkiewicz and Dhonukshe, 2013). Basal land plants posses a single self-renewing apical stem cell (AC) at the growing tip, which governs the entire developmental process (Gifford and Foster, 1989; Hasebe, 2020). The development of three-dimensional body is achieved by sequentially “rotating” the division axis in a helical way with each cell cycle (Harrison et al., 2009; Véron et al., 2021) (Fig. 1a). This rotational division allocates differentiated daughter cells around the AC. Since each daughter cell subsequently gives rise to a portion of the stem and a leaf, rotational division of the AC is indispensable for the organization of the 3D plant body (Moody, 2020).

**Fig. 1.**
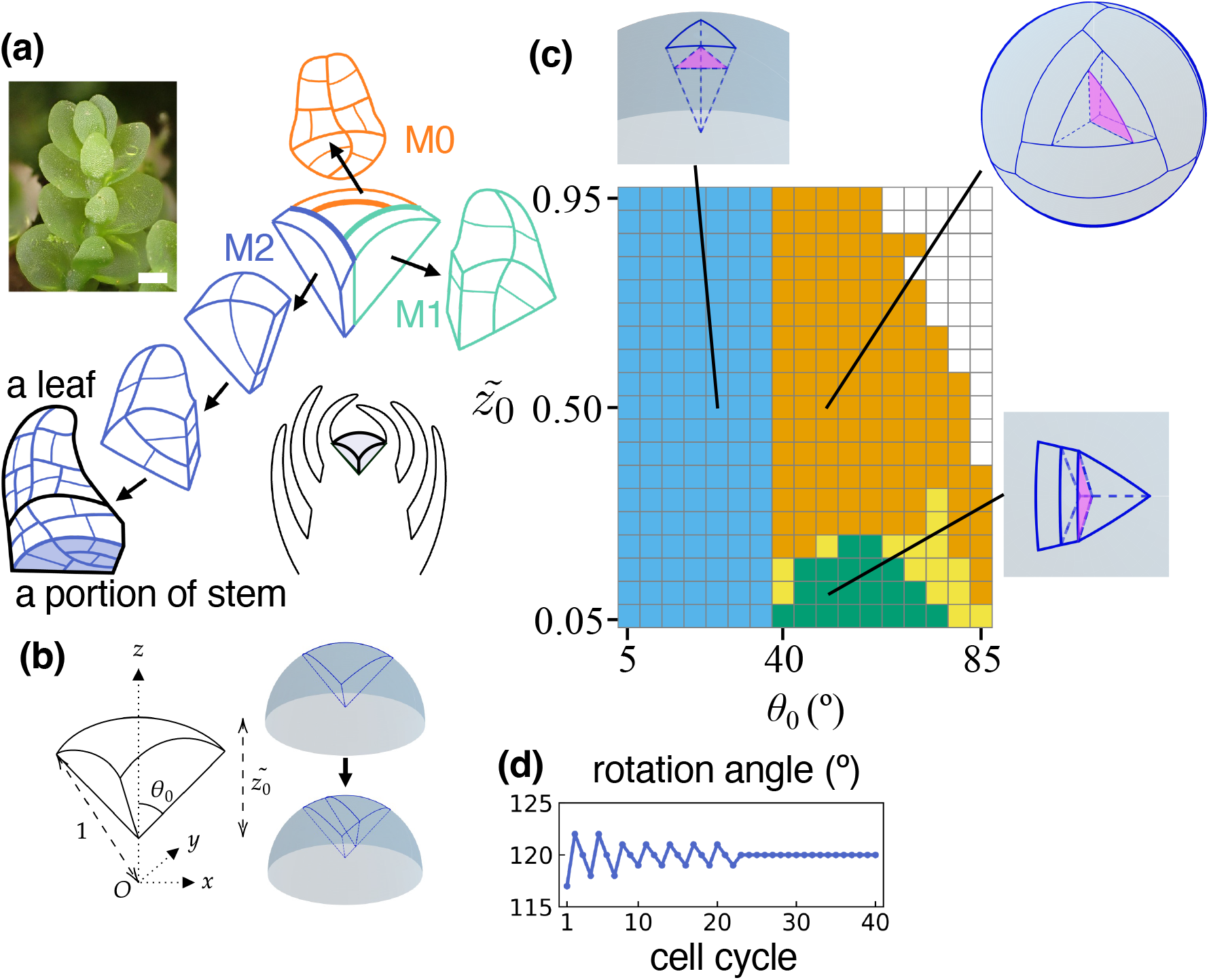
The least area rule governs rotational divisions in the geometrical model. (a) Schematics of the tetrahedral AC, merophyte (M0, M1, M2: labeled in order of the time since cell division from AC), and the corresponding helically symmetric lateral organ arrangement in the leafy liverwort *Haplomitrium mnioides*. The photo was taken at Nichinan, Miyazaki, Japan. Bar: 1 mm. (b) Hemispherical geometric model. The initial shape of tri-symmetric AC is defined by two parameters: polar angle *θ*_*0*_ and dimensionless cell height 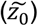 relative to the meristem radius. The division plane is chosen as it passes through the AC centroid and minimizes the area. After division, the daughter with a smaller number of cell vertices is chosen as the next AC, and it isotropically grows to the original cell volume. (c) Phase diagram of simulated division patterns for various *θ*_*0*_ and 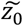. Rotational divisions (orange, top-right inset); periclinal division (blue, top-left inset); parallel divisions (green, right-bottom inset); inconclusive region (yellow). (d) Time course of the rotation angle at *θ*_*0*_ = 50°and 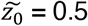.

The 3D shape of the AC and the pattern of its rotational divisions are closely interrelated. The most prevalent shape is the tetrahedron (Fig. 1a; (Hofmeister, 1868; Crandall-Stotler, 1981; Bower, 1889; Gifford and Foster, 1989; Shaw and Renzaglia, 2004; Dolzblasz et al., 2018; Hasebe, 2020; Cruz and Hetherington, 2025). A tetrahedral AC is embedded in the shoot tip, characterized by one free surface and three shared walls with the surrounding differentiated tissues (Fig. 1a). During division, a new wall forms parallel to one of these shared walls, yielding a self-renewing tetrahedral AC and a wedge-shaped differentiated cell. This cell (called merophyte) subsequently differentiates to form a leaf and a portion of the stem (Fig. 1a) (Gifford Jr, 1983; Douin, 1925; Korn, 1993; Harrison et al., 2009). The next division wall in the AC is again oriented parallel to one of the shared walls, but the division axis undergoes a 120° rotation relative to the previous division. The reiteration of such self-renewing, asymmetric cell divisions results in the rotational allocation of differentiated daughters around the AC, simultaneously maintain the AC geometry as a tetrahedron (Fig. 1a). Such structural maintenance may play a role in sustaining the stem cell niche through cell-cell communication, analogous to the mechanisms observed in animal intestinal stem cells (Pentinmikko et al., 2022). Consequently, the rotational angle of the division axis in the AC encodes the rotational symmetry of the organ arrangement, thus determining the overall body plan (Gifford Jr, 1983; Douin, 1925; Korn, 1993; Harrison et al., 2009; Véron et al., 2021; Kamamoto et al., 2021). Understanding of this rotational mechanism paves the way to elucidating how the early land plants acquired their 3D helically-modular body structure. However, the cellular mechanism by which ACs rotate their division axis remains elusive.

One hypothesis for controlling the rotational division is the least area (Errera’s) rule of division plane, which is widely recognized in cell divisions ranging from algae to land plants (Errera, 1886; Berthold, 1886; Thompson, 1942; Cooke and Paolillo Jr, 1980; Miller, 1980; Flanders et al., 1990; Dupuy et al., 2010; Besson and Dumais, 2011; Yoshida et al., 2014; Martinez et al., 2018; Moukhtar et al., 2019; Rong et al., 2022; Ishikawa et al., 2023; Tanaka et al., 2024; Mathew et al., 2025). This rule posits that the division wall minimizes its surface area, analogously to the interface between two soap bubbles within a mother cell. In the early twentieth century, Sir D’Arcy Thompson proposed that the least area rule might govern the arrangement of tetrahedral AC and 120° cells, based on the analogy of his 2D mathematical analysis (Thompson, 1942). The possibility of the least area rule applying to the tetrahedral ACs was also discussed in the observations of the meristems of fern (Lintilhac and Green, 1976) and lycophyte (Karrfalt, 1977). In a more recent study, Couturier and colleagues elegantly showed that the least area rule (i.e., the shortest geodesic path on a surface) in the meristem, approximated as a conical 2D surface, can generate three types of rotation angle (i.e., 180°, 120°, and 90°) seen in ACs (Couturier et al., 2025). Most recently, Cammarata and colleagues proposed a 3D geometrical model incorporating growth fields and repetitive cell divisions, based on the live-imaging of *Physcomitrium patens* AC. Their model quantitatively reproduced the angle of rotational divisions by combining the least area rule with the observed directed displacement of the centroid (potentially the nucleus) of the division plane away from the older existing division walls (Cammarata et al., 2026). Despite the directed centroid displacement concomitant with an asymmetry of the daughter volume ratio in *P. patens* (Cammarata et al., 2026), *the* daughter volume ratio is instead found to be largely symmetric in many other bryophytes, lycophytes, and ferns (Couturier et al., 2025). It thus remains unclear (1) whether and how the least area rule alone achieves the rotational divisions across a broader range of AC 3D geometry (e.g., curvature). An alternative or partially coincident hypothesis is the maximal tension rule. According to this rule, the division wall aligns parallel to the maximal tension axis on the mother cell wall, shown at the shoot apex, root wound healing, and the cambium of *Arabidopsis thaliana*, where anisotropic mechanical stress driven by tissue geometry is dominant (Louveaux and Hamant, 2013; Hoermayer et al., 2024; Höfler et al., 2024). In cases of low stress anisotropy, the maximal tension rule may align closely with the least area rule(Louveaux and Hamant, 2013; Guérin et al., 2016). However, it remains to be elucidated (2) whether and how the maximal tension rule governs the rotational divisions. Finally, identifying (3) the respective functional advantages of the least area and maximal tension rules could help us to understand their roles in 3D tissue organization.

Here, we first developed a simple 3D geometrical model to test whether the least area rule of the volumetrically symmetric division is sufficient to drive steady-state rotational divisions. Next, we developed an elaborate 3D mechanical multicellular model which demonstrated that both the least area rule and the maximal tension rule can independently establish the tetrahedral AC and its helically symmetric division pattern. Our findings reveal the complementary advantages of each rule for developing 3D body organization under plausible cell geometry and mechanics, specifically highlighting how they contribute to geometric self-similarity and developmental robustness.

## 2 Model

### 2.1 3D Geometrical model for hemispherical meristem

The meristem is represented as a hemisphere containing an embedded tetrahedral AC (Fig. 1b). The AC geometry is defined by a spherical free surface and an inner polyhedron, determined by three vertices on the hemispherical surface and one internal vertex (Fig. 1b). To simulate the least area rule, we searched for a division plane that passes through the mother cell centroid and minimizes the interface area. This was achieved by uniformly sampling the normal vector as 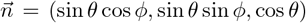, where θ and ϕ are the polar and azimuthal angles, respectively (Fig. S1, top right). The optimal orientation of the least-area division plane was identified 1° increments for both θ and ϕ. After the simulated division, the daughter cell possessing fewer cell vertices is designated as the next AC. The AC then undergoes cell growth until it restores its original volume in an isotropic manner (Fig. S1, right-bottom), following observations in early stages of development (reviewed in (de Keijzer et al., 2021)) and previous models(Kamamoto et al., 2021; Couturier et al., 2025).

#### Rotational angle of division direction

The rotational angle is defined as the angle between the outward normals of successive division planes. This angle is calculated by projecting the normal vectors onto the horizontal (XY) plane and measuring the angular displacement in either a clockwise or counterclockwise direction, consistent with the direction of the rotational divisions.

### 2.2 3D Mechanical multicellular model

To simulate biologically plausible cell deformation, growth and division, we employed the triangular biquadratic spring model (TRBS model) (Delingette, 2008), which has been previously applied to plant multicellular modeling (Bozorg et al., 2014; Bonfanti et al., 2023). In the TRBS model, the cell wall is represented as a mesh of triangular elements (Figs 4a and S3) assuming the hyperelastic material characterized by Young’s modulus Y and Poisson’s ratio σ (supporting text). At each time step, cell wall deformation was computed by minimizing the elastic energy of TRBS (supporting text) under a constant turgor pressure P. Subsequently, cell wall growth was simulated in a strain-based way following the previous study (Bonfanti et al., 2023) (supporting text). To simulate indeterminate growth of the meristem, some topological remeshing operations were introduced along with the cell division (Figs 4a, S3, and supporting text). Cell growth was not considered for merophytes that detached from the AC. All developmental simulations started from a spherical cell with a radius of 10 µm, composed of 162 vertices. We confirmed that the qualitative behavior is robust across a range of parameter values (Y and P; Figs S4 and S5). The simulation was implemented using Julia(Bezanson et al., 2017) and visualized using ParaView(Ahrens et al., 2005).

#### Maximal tension rule

We introduced the maximal tension rule on the curved free surface of the AC using the following procedure: The cell division axis 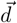 (defined as the unit normal to the upcoming division wall) is determined by identifying the orientation that best fit align with the direction of maximal tension across all free surface triangular elements. For each triangular element *i*, the weighted contribution to the division orientation is evaluated as the absolute inner product 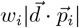 where 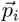 denote the normalized minor eigen vector of the second Piola-Kirchhoff stress tensor. This minor eigenvector is, by definition, perpendicular to both the major eigenvector (the direction of maximal tension) and the face normal of the triangular element. The weight w_*i*_ is defined as the product of the triangle area a_*i*_ and the eigenvalue of the major axis λ_1_ (the intensity of maximal tension), multiplied by the stress anisotropy (λ_1_ − λ_2_)/λ_1_, i.e., w_*i*_ = a_*i*_(λ_1_ − λ_2_). These weighted contributions are summed over all the triangular element constituting the AC free surface for each vector 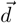. The final division axis is selected as the vector 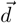 that maximizes this summed objective function.

#### AC determination

Following cell division, the next AC is identified using a criterion similar to that of the geometrical model. To designate the daughter cell with fewer topological vertices as the next AC, we defined “vertices” based on cellular junctions: vertices shared by three cells on free surfaces and those shared by four cells within the inner tissue were counted as the corresponding AC vertices.

#### Sustainability of rotational division against orientation fluctuation

To evaluate the robustness of rotational divisions, we first allowed an initial spherical cell to progress through six cell cycles under the least area rule (e.g., Fig. 3b). Subsequently, we introduced stochastic angular fluctuations to the division axis for the following four cell cycles under either the least area or maximal tension rule. The magnitude of the angular fluctuation was sampled from a von Mises distribution (the circular analogue of the normal distribution) with standard deviations of 2°, 4°, 8°, 16°, and 32° (*n* = 150 independent trials for each value). The axis of this perturbation was randomly selected from a plane perpendicular to the predicted division axis.

After the simulations, we determined whether the AC sustained rotational divisions in the same direction as observed prior to the imposed fluctuation.

## 3 Results

### 3.1 Least area rule of cell division plane reproduced 120° rotational divisions in 3D geometrical model

First, we examined the simplest 3D condition in which an AC is established in a hemispherical meristem to determine whether the least area rule of volumetrically symmetric division plane can achieve the self-renewing rotational divisions as a steady state. We initialized the model with 3D tri-symmetric tetrahedral AC geometries, where the relative height 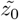 was normalized by the meristem radius (Figs 1b and S1). The least area division plane was numerically determined by comprehensively searching the orientations passing through the AC centroid in 3D (Fig. S2; Method). After cell division, the tetrahedral daughter was designated as the next AC (Fig. 1b), which subsequently underwent isotropic growth until it regained its original volume (Fig. S1; Method). We found that consecutive rotational divisions emerge across a broader parameter space of the initial geometry (e.g., the polar angle ≥ 40° and the cell height > 0.2 in Fig. 1b, c; Video S1). The rotational angle between successive division planes (roughly equivalent to the divergence angle; (Kamamoto et al., 2021)) was approximately 120° (Fig. 1d), consistent with those observed in typical leafy liverworts and ferns. Notably, these rotational divisions disappeared below a threshold of the height-to-meristem radius ratio 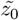, which correlates with the curvature of the AC free surface (Fig. 1b). These results demonstrate that the least area rule is sufficient to accurately reproduce the rotational divisions, provided the AC free surface possesses adequate curvature.

### 3.2 Rotational division is ensured by rotation of geometric proportion of AC

Given that the least area division only depends on the 3D geometry (shape) of AC, we hypothesized that each division event rotates the geometric proportion of AC. To test this theory, we quantified the geometric similarity of the AC outline proportions before and after division by calculating the similarity of each arc and edge length of the AC outline (Fig. 2a, left). At high curvature, the geometric similarity was maximized when the AC geometry after cell division is compared with the geometry before cell division under 120° rotation. This rotational self-similarity supports our theory of rotation of the geometric proportion (Fig. 2a, 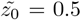). In contrast, at low curvature, the geometric similarity is maximized with no rotation (Fig. 2a, 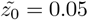), which is consistent with the parallel divisions (Fig. 1c). These results indicate that the least area division of the highly curved ACs ensures the rotation of the AC geometric proportion.

**Fig. 2.**
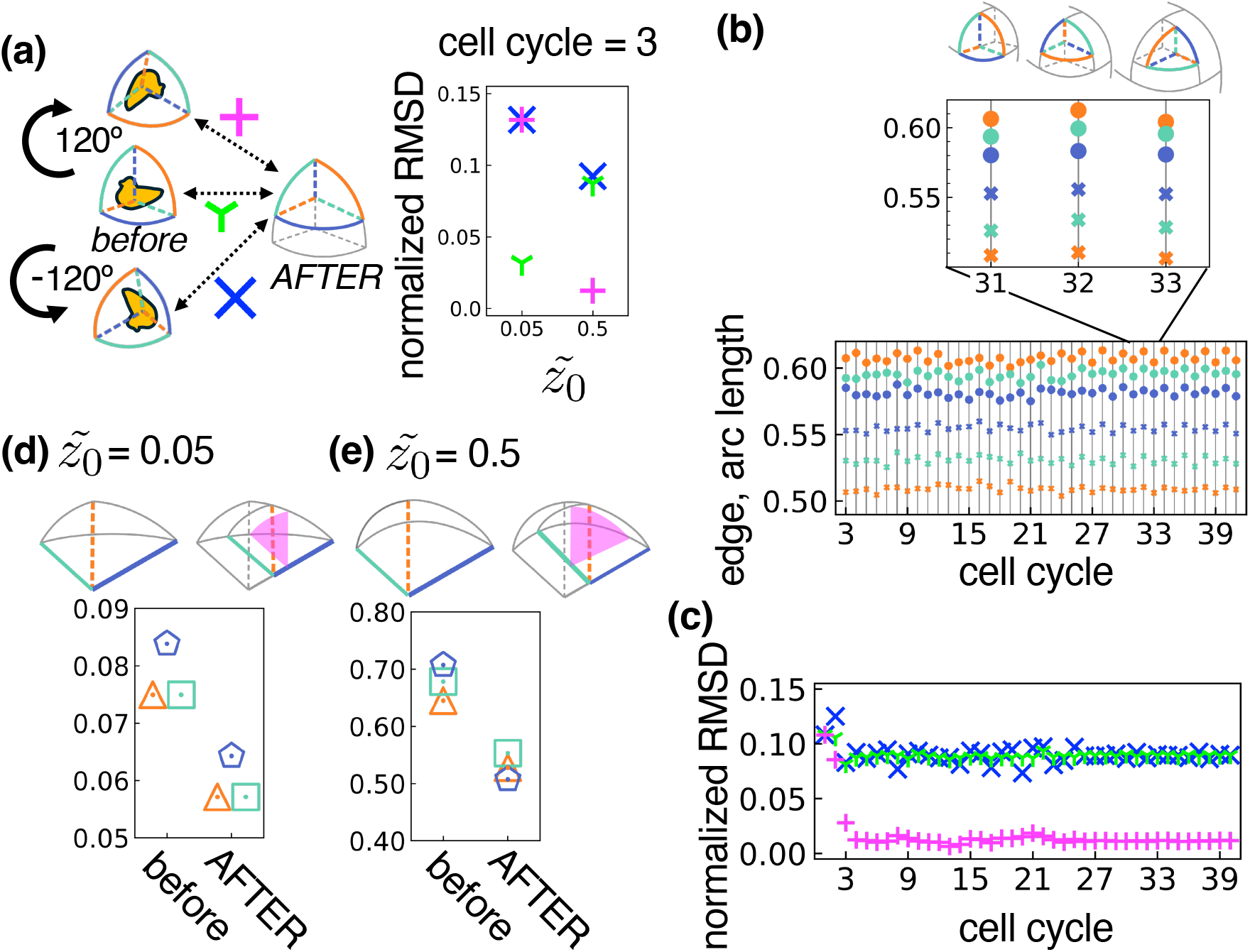
Geometrical analysis of rotational divisions. (a) The root-mean-square deviation (RMSD) between the AC outlines before and after cell division was calculated by 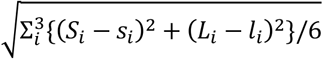, where *S*_*i*_ and *L*_*i*_ denote the arc length and edge length, respectively, while *s*_*i*_ and *l*_*i*_ denote corresponding lengths after division and growth. At two different curvatures 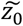 (right panel), RMSD was measured for three configurations (left panel): parallel (green Y-markers), rotated (magenta plus signs), and reversely rotated (blue crosses), then normalized by the characteristic length scale of AC (AC volume)^(1/3). (b, c) Time course of lengths of AC outline edges and arcs (b) and RMSD (c) at 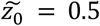. The arcs and the edges are colored according to the age of the adjacent and opposite merophytes, respectively (orange: M0, green: M1, and blue: M2). (d, e) Comparison of AC outline edge lengths before and after the division at 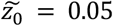 (d) and 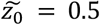 (e). *θ*_*0*_ is set to 50°.

To determine whether the curvature-mediated rotation of geometric proportion during each cell cycle leads to consecutive rotational divisions in a constant direction, we tracked the time course of AC geometry using edge and arc lengths (Fig. 2b) at high curvature of AC 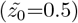. Each arc length monotonically decreased over successive cell cycles (e.g., orange → green → blue circles in Fig. 2b inset). This pattern revealed that the arc shared with the youngest and oldest differentiated cells is always the longest and shortest, respectively (Fig. 2b). This exemplifies the consecutive 120° rotation of the AC proportion for each cell cycle. Accordingly, the geometric self-similarity is always maximized with 120° rotation in a fixed direction (Fig. 2c, magenta plus signs).

Finally, to understand the mechanism underlying the rotation of the AC geometric proportion at high surface curvature, we analyzed how the insertion of a least area division wall affected the edge lengths of the AC outline (Fig. 2d, e). At both low and high curvatures, the division wall intersects the longest edge of the AC outline (Fig. 2d, e top-left, blue lines). However, at low curvature, this edge remained longer than the newly formed edges (Fig. 2d top-right, green and orange lines). Since the subsequent division wall again intersected this longest edge, the division axis is likely to be the same as the previous one (Fig. 2d top-right, magenta). In contrast, at high curvature, a newly formed edge (Fig. 2e top-right, green line) increased in length due to the highly curved arc on the free surface (Fig. 2e, top grey curve), and it exceeds the previously longest edge (blue line). This process reordered the edge lengths (from Fig. 2e left, blue > green > orange to Fig. 2e right, green > orange > blue), thereby resulting in the rotation of the AC’s geometric proportions (Fig. 2e, top right). Taken together, these results reveal that the least area division-driven AC rotational self-similarity ensures the consecutive rotational divisions in 3D.

### 3.3 Least area rule under plausible cell mechanics establishes tetrahedral AC and the rotational divisions

We next investigated whether the rotational divisions driven by the least area rule could function under plausible cell growth and deformation mechanics. To this end, we developed an AC-driven developmental model based on the continuum mechanics-consistent mechanical model (TRBS model, (Delingette, 2008; Bozorg et al., 2014; Bonfanti et al., 2023); Fig. S3). We introduced cell division (Fig. 3a; supporting text) and indeterminate meristem growth using remeshing algorithms (Fig. S3; supporting text). Both the AC and the adjacent differentiated cells grew through uniform turgor pressure and cell wall extension (Methods; (Bonfanti et al., 2023)). The AC was specifically divided when it reached a threshold cell volume. In each cell division, the daughter cell with fewer vertices was designated as the next AC. Starting the simulation from a single spherical cell, three successive morphologically symmetric divisions spontaneously yielded a tetrahedral AC (Fig. 3b top row; Video S2). Subsequent divisions involved inserting the division wall parallel to the oldest division wall surrounding the AC, resulting in consecutive rotational divisions (Fig. 3b bottom row, c; Video S2). The tetrahedral AC and consecutive rotational divisions were reproduced within a certain range of typical cell wall stiffness and turgor pressure values (Fig. S4, Method). Throughout the simulation time courses, the AC geometry maintained the rotational self-similarity with 120° (Fig. 3d, e), consistent with the results from the geometrical model (Fig. 2a-c). These results indicate that the establishment of a tetrahedral AC and the maintenance of 120° rotational divisions are natural consequences of the interplay between least area division and turgor pressure-driven 3D cell growth.

**Fig. 3.**
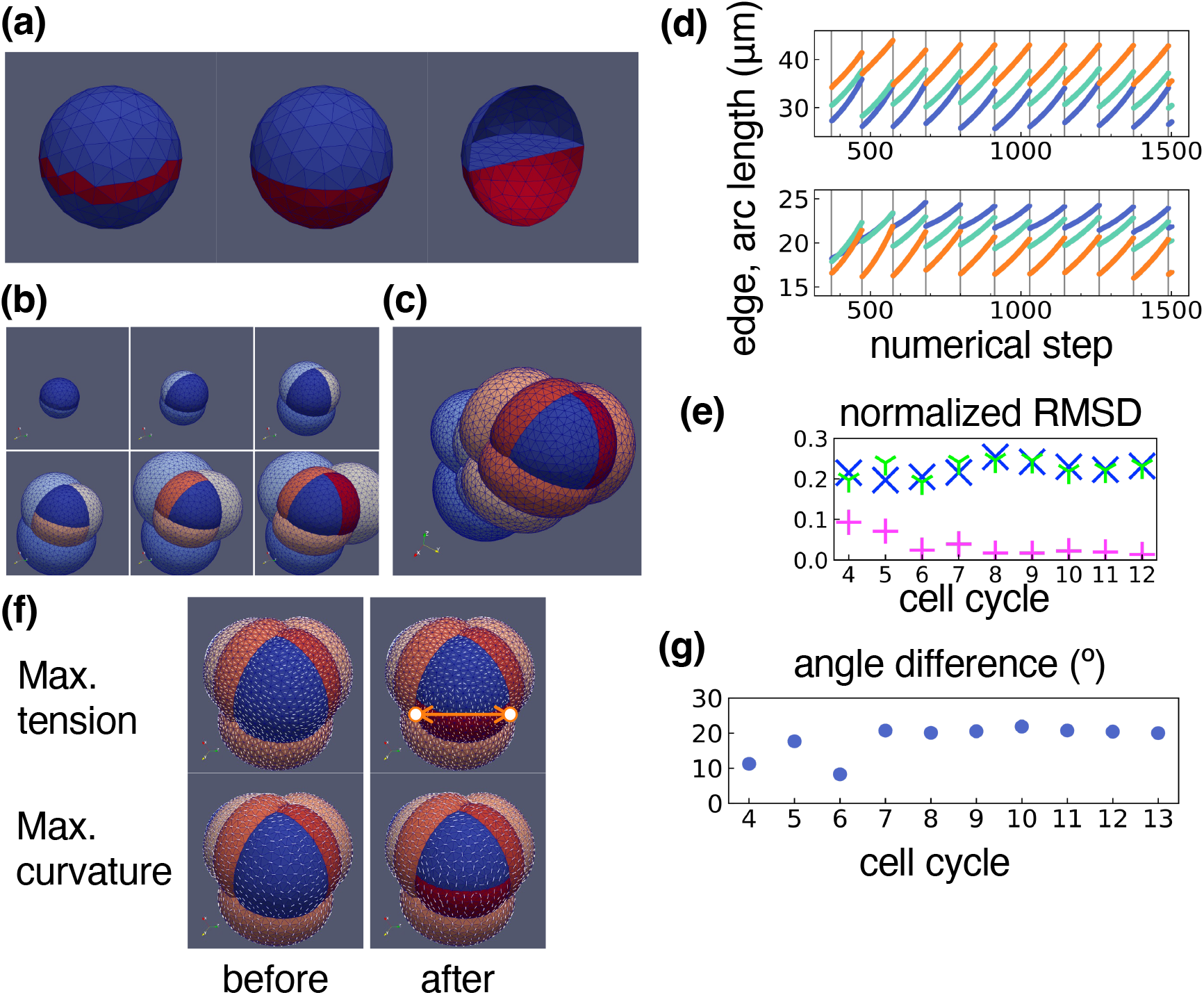
Rotational divisions based on the least area rule in the mechanical model. (a) Outline method of a cell division: identification of the triangle elements intersected by the division plane (left) and subsequent remeshing of these elements (center). The sectional view is shown on the right. The cell wall is represented by triangular elements. (b) Snapshots of repetitive least area divisions of the AC (colored in deep blue). Three morphologically symmetric cell divisions (top row) and subsequent asymmetric-rotational divisions (bottom row). (c) Snapshot after 12 cell divisions. (d, e) Time course of AC outline lengths (edges and arcs) (d) and RMSD (e). Colors and markers are identical to those used in Fig. 2b and 2c, respectively. (f) The maximal tension axis (top row) and maximal curvature axis (bottom) of each triangle element just before (left column) and after (right) the least area division. (g) Angle difference between the mean maximal tension axis and the division plane tangent. The tangent is defined by the vector connecting the AC-M0-M1 junction to AC-M0-M2 junction on the free surface (orange two-way arrow in (f)). The initial radius of the spherical cell is 10 μm. Turgor pressure *P* = 0.2 MPa, Young’s modulus *Y* = 100 MPa, and Poisson’s ratio *σ* = 0.2.

### 3.4 Maximal tension rule reproduced 120° rotational divisions ensured by the history of division wall formation

We found that immediately prior to rotational divisions, both the maximal tension patterns and the maximal curvatures on the AC free surface tended to align parallel to the subsequent division wall, in accordance with the maximal tension rule (Fig. 3f, g). Therefore, we incorporated the maximal tension rule into our mechanical model (Methods), replacing the least area rule. While this rule did not yield the rotational divisions when applied to both the AC free surface and the AC-merophyte shared walls, it successfully reproduced rotational divisions when applied exclusively to the AC free surface (Fig. 4a, S5; Video S3). Unlike simulations governed by the least area rule, rotational self-similarity of the AC geometry is maintained for several cell cycles at most (Fig. 4b, c). Nevertheless, the rotational divisions persisted.

**Fig. 4.**
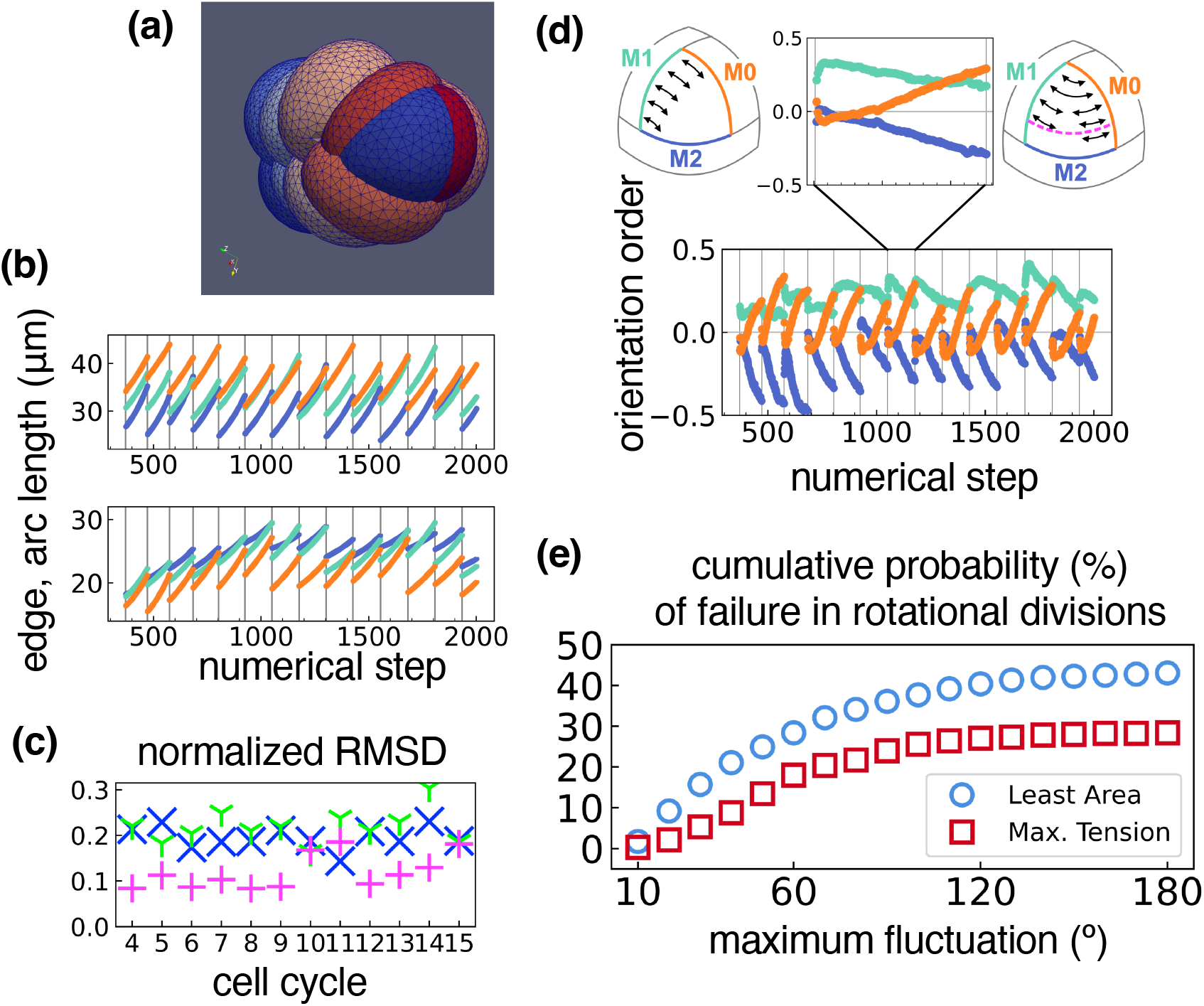
Rotational divisions based on the maximal tension rule. (a) Snapshot after 12 cell divisions. (b, c) Time course of AC outline lengths (edges and arcs) (b) and RMSD (c). Colors and markers in (b, c) are identical to those used in Fig. 2b and 2c, respectively. (d) Bottom: Orientation order parameter of the maximal tension axes on the AC free surface against each orthogonal axis of the AC-merophyte shared walls (AC-M0: orange, AC-M1: green, and AC-M2: blue). The parameter is defined by ∑_*i*_(1/2)*a*_*i*_(3cos^2^(*θ*_*i*_) − 1)/ ∑*a*_*i*_, where *a*_*i*_ denotes the area of triangle element *i*, and *θ*_*i*_ is the angle between the shared wall normal and the maximal tension axis. Top: Representative pattern of the maximal tension (black two-way arrows) on the AC free surface. The future division wall is indicated by the magenta dashed line. (e) Cumulative probability of rotational division failure under fluctuations in the division axis, comparing the least area and the maximal tension rule. The parameters are consistent with those shown in Fig. 3. *p-value < 0.001 (Kolmogorov–Smirnov test).

To identify the mechanical properties that ensure consecutive rotational divisions in the maximal tension rule without requiring the AC rotational self-similarity, we analyzed how each division affected the maximal tension pattern on the free surface (Fig. 4d). Immediately after each cell division, the maximal tension axes aligned primarily with the AC-M1 shared wall axis (represented by the normal vector). This is followed by alignment with the AC-M0 (the most recently formed division wall) and the AC-M2 shared wall (Fig. 4d, bottom, inset and top left). As the cell cycle progressed, the maximal tension axes rotated to align with the AC-M0 shared wall axis, while maintaining alignment with the AC-M1 shared wall axis (Fig. 4d bottom, inset and top right). This indicated that the newly formed division wall (i.e., AC-M0 shared wall) pulled the AC free surface inward, generating the tension pattern aligned with the AC-M0 shared wall axis. As these tension patterns were gradually overwritten by the subsequent formation of division walls, by the end of the cell cycle, the tension axes aligned with both the AC-M0 and AC-M1 shared wall axes, but remained perpendicular to the AC-M2 shared wall axis (Fig. 4d top right). These results suggested that the history of the division wall formation through transient tensional patterns ensures consecutive rotational divisions even when the AC self-similarity is not strictly maintained.

### 3.5 Maximal tension criterion more robustly maintains the rotational divisions against fluctuations

The emergence of rotational divisions by both the least area and the maximal tension rules, but based on different mechanisms (Figs 3b-e and 4a-d), prompted us to identify the distinct advantages of each rule in 3D tissue organization. While the least area rule is advantageous for maintaining the rotational self-similarity (Fig. 3e and 4c), we examined whether the maximal tension rule confers greater robustness to rotational divisions, facilitated by the history of division wall formation. We thus compared the robustness of the persistence of rotational divisions between the two rules in the presence of stochastic angular fluctuations (Methods). At small fluctuations (< 10° in Fig. 4e) for four cell cycles, both rules robustly sustained rotational divisions in the same direction as established prior to the introduction of fluctuation. As the intensity of fluctuations increased, the difference in the probability of sustained rotational divisions between the two rules widened significantly, demonstrating that the maximal tension rule confers greater robustness against orientational fluctuation compared to the least area rule (Fig. 4e). These results suggested that the maximal tension rule favors rotational persistence against orientational stochasticity over the strict maintenance of rotational self-similarity.

## 4 Discussion

Building on the seminal theoretical study of Couturier and colleagues that meristem angle controls the rotational divisions on a conical 2D surface (Couturier et al., 2025), here we have presented a 3D geometrical model of the meristem. In our model, the AC is represented as a volumetric entity defined by three internal walls meeting at a shared vertex and a curved surface. By incorporating isotropic volumetric growth and symmetric cell division following the least area rule into this geometrical model, we showed that, consistent with the meristem angle (Couturier et al., 2025), the curvature of AC surface controls the rotational division in 3D, through the rotation of the geometric proportion of the AC (Fig. 2). Such 3D self-replication through rotation of AC geometric proportion seemingly forms a basis for self-similar growth of plant bodies and modular arrangement of merophytes with helical symmetry. Furthermore, our mechanical model confirmed that such 3D self-replication can be robustly achieved and maintained even under biologically plausible growth mechanics (Figs. 3b-e, S4). Therefore, our theoretical results reveal a mechanistic chain wherein the least area rule and high surface curvature ensure 3D self-replication leading to the rotational divisions. These factors are testable *in vivo*, by applying the presented or earlier methods (Martinez et al., 2018; Moukhtar et al., 2019; Ishikawa et al., 2023; Tezuka et al., 2025; Cammarata et al., 2026) to 3D volumetric data. Specifically, the least area rule can be verified by determining whether the area-minimizing plane passing through the cell centroid (Fig. S2) consistently coincides with the actual division plane. The 3D self-replication could also be confirmed if the presented RMSD is minimized between two cell cycles of AC after a 120° rotation in a constant direction (as seen in Fig. 2a, c). The consecutive rotational divisions driven by symmetric daughter’s volumes (Figs. 1b-c and 3b-c) are observed in bryophytes and ferns Acs (Couturier et al., 2025). Specifically, our model does not require volumetrically asymmetric divisions due to directed centroid displacement, reported in *P. patens* (Cammarata et al., 2026), for rotational division in a fixed direction. Investigating these geometric factors in bryophytes and ancestral algae in future experiments will help elucidate the design principles behind the evolutionary transition from 1D or 2D cell arrangements to complex 3D body symmetry during land plant colonization.

Moreover, our mechanical model (Fig. 3f, g, and 4a) revealed that the spatial distribution and temporal shifts of the maximal tension pattern on the AC surface drives rotational divisions. Under the maximal tension rule, rotational divisions are sustained by the iterative deformation of the AC surface induced by the formation of successive division walls (Fig. 4d). Mechanically, the surface region adjacent to the fringe of a new division wall is pulled inward mechanically, resulting in the emergence of locally high curvature and tension towards the division wall (Figs 3f and 4d). Therefore, the curvature pattern on the surface could serve as a proxy for the maximal tension distribution, helping to predict the division plane *in vivo*. This tension pattern is partially overwritten by subsequent division wall formation (Fig. 4d). Thus, once an AC with three shared walls is established, the integrated tension pattern shifts toward the more recently formed walls (the AC-M0 and AC-M1 shared walls) (Fig. 4d). Consequently, the next division wall is consistently oriented parallel to the oldest (AC-M2) shared wall, resulting in persistent rotational divisions. Such dynamic shifts in tension have been suggested to influence the cell division orientation in angiosperms (Colin et al., 2020). It would therefore be worthwhile to examine in the future whether such dynamic couplings between cell geometry and tension patterns further govern the rotational division of AC.

Importantly, the rotational divisions mediated by the history of division wall formation exhibit greater robustness against fluctuations in the division axis than those mediated by the geometric rotational self-similarity under the least area rule (Fig. 4e). This enhanced robustness arises because such fluctuation little affects the tension pattern on the surface near the fringe of the newly formed division wall. In contrast, similar fluctuation can trigger aberrant rotation or non-rotation of the geometric proportion of AC, subsequently leading to the misorientation of the least area division plane. Considering that redundant processes regulate cell division orientation *in vivo* (Uyehara and Rasmussen, 2023), *the maximal tension and the least area* rules may play complementary roles: the former could govern the initial spindle orientation to ensure directional persistence, while the latter refines the exact orientation by minimizing the division wall area. Through this cooperation, plants may achieve both developmental robustness and geometric precision in 3D tissue organization.

The cell-based architecture of our mechanical model (Figs 3a and S3) allows for the integration of diverse cytological regulators of cell division, growth, and differentiation. These include gene regulatory networks, PIN-mediated polar transport of auxin (Yoshida et al., 2014), plasmodesmata (Abitbol-Spangaro et al., 2025), microtubules (Bozorg et al., 2014), and nuclear displacement (Robinson et al., 2011; Cammarata et al., 2026). The detailed 3D cell geometry, repetitive cell divisions, and mechanical stress patterns (presented in Figs 3b, c, f, and 4a) provide a robust framework to simulate the morphogenesis of multicellular and multilayered plant bodies and organs that originate from a single or a small number of cells. One promising area for future research is explaining alternative rotation angles (e.g., 180°), which are observed in bryophytes and ferns, and were predicted to occur at steep meristem angles in the conical 2D surface model (Couturier et al., 2025). If the least area rule in 3D can reproduce a 180° rotation angle, the required surface curvature of the meristem would likely be steeper than both the spherical surface (Fig. 1b) and some deviation from the spherical surface due to cell-cell adjacent topology and geometry of both AC and surrounding merophytes (Figs 3b, c, and S4). Such steep curvatures of AC may be driven by tip growth or cortical microtubules-driven anisotropic diffuse growth (Ren et al., 2017) of AC. Alternatively, the maximum tension rule might account for such rotation angles without a visible steep surface in the AC. Instead, this could due to anisotropic stress states imposed by surrounding merophytes, potentially arising from cortical microtubules-mediated tissue-flattening (Zhao et al., 2020) or dorsoventrality (Shaw and Renzaglia, 2004; Plackett et al., 2015; Cruz and Hetherington, 2025). Thus, extending the present model to incorporate growth anisotropy (e.g., tip growth (Dumais et al., 2003; Kang et al., 2024) and/or ordering the cortical microtubules (Bozorg et al., 2014; Zhao et al., 2020)) will likely reveal how diverse rotational angles are modulated across land plant evolution.

Another future extension is the application of our model to explain rotational divisions in the root apical meristem of lycophytes and ferns. In addition to anticlinal rotational divisions relative to the distal cell wall, periclinal divisions parallel to the distal wall also occur to produce the root cap (Imaichi and Kato, 1989; Aragón-Raygoza et al., 2020). Since the periclinal division plane exclusively intersects the sidewalls, the cue for its orientation likely follows the tension distribution along these sidewalls. Incorporating this sidewall tension pattern would clarify the relative contribution of the sidewalls and the distal wall to the mechanical switching between division pattern. such an analysis could, in turn, provide insights into the developmental transition from aerial to subterranean organs.

Other future extension is to account for early embryogenesis, in which the tetrahedral cells typically originate from a single spherical cell (Fig. 3b) in angiosperms (Gifford and Foster, 1989; Dresselhaus and Jürgens, 2021). While the emergence of such tetrahedral cells has been shown to be regulated by the least area rule in the Arabidopsis embryo(Yoshida et al., 2014; Moukhtar et al., 2019), it has also been reported that the division planes deviate from the least area plane in the rice embryo (Tezuka et al., 2025). Our finding that tetrahedral cells can emerge through the maximal tension divisions (Fig. 4) provides a potential mechanistic explanation for such deviations. Incorporating the surrounding tissue geometry of various angiosperm embryos into our model will enable us to test whether the maximal tension or least area rule predominantly accounts for the early embryogenesis. These future extensions of the present model could predict the potential regulatory factors underlying the conservation and diversification of plant morphogenesis driven by a small number of initial cells.

## 5 Acknowledgements

We thank Masaki Shimamura (Hiroshima University, Japan), Satoru Tsugawa (Akita Pref. University, Japan), and Minako Ueda (Tohoku University, Japan) for valuable discussions and suggestions. This work is supported by the Japan Science and Technology Agency (CREST [JPMJCR2121 to N.K. and K.F.]).

## 6 Author contributions

NK and KF conceived and designed the research. NK developed the mathematical models, wrote the code, and performed the simulations. NK and KF wrote the manuscript.

