## Supporting Text for "3D geometry and mechanics of a single apical stem cell ensure helically symmetric plant body in multicellular models"

February 2026

### 1 Geometrical model

#### 1.1 AC growth

Following each cell division, the AC grows isotropically to restore the original cell volume. To maintain the spherical geometry of the AC surface, the growth ratio is determined according to the following procedure. First, the inner polyhedral portion (i.e., all the vertices of the AC) is isotropically expanded by a provisional ratio larger than 1. Second, this polyhedron is translated downward along the mean unit normal vector $\boldsymbol{n}$ of the AC free surface to ensure that the vertices originally located on the surface prior to expansion again intersect the spherical surface. This descent distance $d$ is derived from the intersection condition:

$$(\boldsymbol{x} - d\boldsymbol{n})^2 = 1 \tag{1}$$

where $\boldsymbol{x}$ represents the one of the surface vertices. Solving this equation for $d$ yields:

$$d = \boldsymbol{x} \cdot \boldsymbol{n} - \sqrt{(\boldsymbol{x} \cdot \boldsymbol{n})^2 - (\boldsymbol{x} \cdot \boldsymbol{x})^2 + 1}. \tag{2}$$

After these procedures, the total AC volume is calculated. The appropriate growth ratio is numerically searched until the total AC volume converges to the target volume.

#### 1.2 Merophyte growth

For visualization purposes only, merophyte vertices located on the hemispherical surface were translated toward the periphery. This displacement occurred along the arc defined by the positional vector of the vertex and the normalized positional vector of the AC centroid. The angular displacement along this arc per each AC cell cycle was defined as $(2/3)V^{1/3}\Delta\theta$, where $V$ and $\Delta\theta$ are the original AC volume and the angle between the two vectors, respectively.

### 2 Mechanical model

The elastic strain energy of the St. Venant-Kirchhoff hyperelastic material is expressed as:

$$W_\Omega = \int_\Omega \left( \frac{\lambda}{2} (tr\boldsymbol{E})^2 + \mu \boldsymbol{E} : \boldsymbol{E} \right) d\Omega \tag{3}$$

where $\boldsymbol{E}$ is the Green-Lagrange strain tensor and : denotes the tensor dot product. $\lambda$ and $\mu$ are Lamé constants:

$$\lambda = \frac{Y\sigma}{1 - \sigma^2}, \ \mu = \frac{Y}{2(1 + \sigma)}. \tag{4}$$

In the TRBS formulation, the elastic energy of each triangle element is expressed as (Delingette, 2008):

$$W_{TRBS}(T_p) = \sum_i \frac{k_i^{T_p}}{4}\left(l_i^2 - L_i^2\right)^2 + \sum_{i,j} \frac{c_{i,j}^{T_p}}{2}\left(l_i^2 - L_i^2\right)\left(l_j^2 - L_j^2\right). \tag{5}$$

Here, effective spring constants $k_i^{T_p}$ and $c_{i,j^{T_p}}$ are defined as (Delingette, 2008):

$$k_i^{T_p} = \frac{2\cot^2\alpha_i(\lambda+\mu)+\mu}{16A_p} \quad \text{and} \quad c_{i,j}^{T_p} = \frac{2\cot\alpha_i \cot a_j(\lambda+\mu)-\mu}{16A_p}. \tag{6}$$

$L_j$ and $l_j$ denote the reference and current edge length of the $j$-th edge of the triangle, respectively. $A_p$ is the area of the reference triangle, and $\alpha_j$ is the interior angle of the reference triangle opposite to the $j$-th edge.

At each time step, cell wall deformation is computed by minimizing the elastic energy under constant turgor $P$ using the L-BFGS algorithm (Liu and Nocedal, 1989). Cell wall growth is modeled as the irreversible extension of reference edge length, following previous study (Bassel et al., 2014; Bonfanti et al., 2023) as:

$$L_j^{new} = L_j + \Delta t\Phi\left[\frac{l_j}{L_j} - 1\right]_+ L_j \tag{7}$$

where $\Phi$ denotes the extensibility and $[]_+$ is the ramp function (Fig. S3, right bottom). To ensure computational efficiency and minimize the frequency of remeshing operations, the efective growth ratio $\Delta t\Phi$ was set to $0.04 * Y/P$. This parameterization maintained the cell cycle at approximately 100 growth steps per AC division.

#### 2.1 Remeshing

To maintain numerical accuracy during cell growth and to prevent unexpected geometrical and/or topological errors during cell division, we implemented three remeshing operations based on (Okuda et al., 2022; Khan et al., 2022).

**Vertex centering:** To enhance homogeneity in triangle shapes, a centering operation is performed by moving each vertex toward the centroid of its connecting triangles. To ensure volume conservation, the vertex is translated along the mean outward normal vector to a position that maintains the cell volume. The new position of the target vertex $\boldsymbol{x}_{new}$ is expressed as:

$$\boldsymbol{x}_{new} = \boldsymbol{c} + h\boldsymbol{n}. \tag{8}$$

$\boldsymbol{n}$ is the unit mean outward normal vector of the connected triangles. The centroid $\boldsymbol{c}$ is defined as:

$$\boldsymbol{c} = \frac{\sum_i a_i \boldsymbol{n}_i}{\sum_i a_i}. \tag{9}$$

where $a_i$ is the area of the triangle $i$ sharing the vertex. The scalar displacement $h$ is determined by minimizing the potential function:

$$U = (V - V_0)^2 \tag{10}$$

where $V$ and $V_0$ are current and target cell volume, respectively.

To minimize the effect of the centering operation on the mechanical state of the surrounding mesh, we adopted Bozorg's formulation (Bozorg et al. (2016)) to calculate the reference edge lengths after the centering as:

$$L = \left[\sum_i (1 - 2\lambda_i)|\boldsymbol{t} \cdot \boldsymbol{s}_i|^2\right]^{1/2} \tag{11}$$

where $\lambda_i$ and $\boldsymbol{s}_i$ are the eigenvalues and eigenvectors of the mean strain tensor, respectively, and $\boldsymbol{t}$ is the edge tangent vector.

**Edge splitting:** Edge splitting is performed to increase the number of triangle elements and maintain the geometrical accuracy. When the area of two adjacent triangles exceeds a predefined threshold value, a new vertex is inserted at the midpoint of the shared edge, dividing each triangle. This operation is applied for both manifold and non-manifold (tricellular or bicellular and cortical junction) edges. In manifold edges, the new reference edge lengths are determined in the same way as in the centering. In non-manifold edges, each reference length is set as $(1-\epsilon)l$, where $l$ is the current length, and $\epsilon$ is a small constant. To prevent local perturbation in the stress state, splitting is suppressed within the neighbors of triangles that have already been split during the same growth step.

**Edge flipping:** Edge flipping is performed to regularize the triangle geometry and reduce the "narrow" (high aspect ratio) triangles. To prevent destructive topology changes, this operation is restricted by the following conditions:

1. The target edge must be a manifold edge.
2. The number of edges connecting at each vertex of the target edge (i.e. *degree*) must be greater than three.
3. The edge length must exceed a predefined threshold value.
4. After flipping, the interior angles adjacent to the new edge must exceed a specific threshold angle.

The last condition is necessary to prevent the formation of locally concave geometries.

#### 2.2 Cell division

Given the cell division axis following the least area or the maximal tension rules, cell division is operated as follows. First, the triangle elements intersected by the future division plane, which passes through the cell centroid and is perpendicular to the division axis, are identified (Fig. 3a left). Subsequently, the shortest closed loop is identified along the edges of these elements. Second, the vertices constituting the shortest loop are projected to the future division plane so that these vertices and edges become the perimeter of the future division plane (analogy of the cortical division zone). To maintain the mother cell volume and minimize perturbations to the mechanical state, these projections are performed iteratively (e.g., 30 steps) in conjunction with centering operations. Finally, new vertices constituting the division plane are generated by the bubble-mesh method (Shimada and Gossard, 1995) and integrated into the triangle mesh by Delaunay's triangulation (Fig. 3a center and right). The reference edge lengths of the newly generated meshes were set to 0.99 times the current lengths to ensure numerical stability.
